# Can combined compost and biochar application improve the quality of a highly weathered coastal savanna soil?

**DOI:** 10.1101/2020.07.24.219279

**Authors:** Kwame Agyei Frimpong, Emmanuel Abban-Baidoo, Bernd Marschner

## Abstract

Soil fertility decline represents a major constraint to crop productivity in sub-Saharan Africa. Many studies have shown that addition of biochar or compost can effectively improve soil quality. Biochar produced from crop residues are often N-poor but rich in stable C while poultry manure composts, which is often rich in nutrients including N decomposes rapidly under high rainfall and temperature conditions. Combined biochar and compost application can compensate for the shortcomings of each other such that their interactive effect is likely to improve soil quality. A 30-days incubation experiment was carried out on a Haplic acrisol amended with corn cob biochar, rice husk biochar, coconut husk biochar, poultry manure compost and composted rice husk or corn cob biochar to examine the effect of compost and biochar, applied singly, in combination or as co-compost on basal soil respiration, and soil quality indicators such as soil pH; soil microbial carbon; cation exchange capacity; total organic carbon, total nitrogen and available nitrogen concentration. The results showed that addition of the different amendments increased soil pH compared with the untreated control with the combined corn cob and rice biochar and compost treatments recording the highest pH values. Basal respiration following sole compost, composted biochar and combined biochar and compost application were significantly greater than the sole biochar and the control treatments. TOC increased by 37% in the sole compost treatment to 117.3% in the combined corn cob biochar and compost treatment, respectively. MBC increased by 132.2% in the combined rice husk biochar and compost treatment and by 247% in the sole compost treatment compared to the control. The study has demonstrated the potential of compost, biochar and especially composted biochar to enhance soil quality, C stabilization and reduce soil C loss through basal respiration.

## Introduction

Improving soil quality is critical to increasing crop yields, combating rural poverty and reversing natural resource degradation in sub-Saharan Africa (SSA) [1]. Biochar, a highly stable, aromatic carbon by-product obtained from the pyrolysis of organic material at relatively low temperatures (<700 °C) in the presence of low or no oxygen, has been shown to be an effective amendment to improve soil quality [2]. Biochar is not biologically inert but follows a biphasic mineralization pattern where the more labile compounds are rapidly mineralized first, after which the recalcitrant carbon degrades more slowly [3]. Consumption of labile compounds from biochar decomposition changes soil physicochemical characteristics [2] and soil biological parameters including basal soil respiration and microbial biomass [4,5]. Although, the rate of biochar decomposition is slower than other soil carbon pools, it can provide similar soil ecological services including nutrient and water retention as other organic materials [6]. Kolb et al. [7] reported that the specific surface area of biochar, which is approximately 200 to 400 m^2^ g^-1^, is comparable to that of the soil clay fraction. Thus, biochar addition can increase the specific surface area of soil [8], increase water and air availability and stimulate microbial activity for enhanced soil quality.

Soil acidity is a major constraint to crop production in many areas of tropical Africa. Liming is the conventional option for soil acidity mitigation, but lime is costly and scarcely available to smallholder farmers in SSA hence its adoption is very low. Van Zweiten et al. [9] found that biochar added to soil exerts a liming effect, which can promote nutrient release from native soil nutrients pools. According to Oguntunde et al. [10], the basic metallic ions contents of the ash component of the biochar applied can decrease soil acidity and create a growth-stimulating effect, especially in soils with low fertility [11]. Biochar application has potential to promote soil C and N retention, increase cation exchange capacity and reduce anthropogenic greenhouse (CO_2_, N_2_O and CH_4_) emissions [12,13].

Compost produced from aerobic decomposition of organic materials has been shown to have significant effects on soil physicochemical properties [14]. For instance composts produced from crop residues, manures and other biomass waste products can increase soil nutrient content [15]. Thus, application of composts is a commonly recommended practice that can introduce nutrients to soil and improve yields in low-input farming systems in SSA. However, under the high temperature and rainfall conditions prevailing in SSA, composts are rapidly mineralized after soil application within months or a few years [16] and the nutrients released may be subjected to leaching or gaseous losses.

Many biochars are N-poor but rich in stable C [13] while most compost are often rich in nutrients including N. Combined biochar and compost application can compensate for the shortcomings of each other such that their interactive effect is likely to improve soil quality. Liu et al. [17] observed a synergistic positive effect of compost and biochar mixtures on soil organic-matter content, nutrients concentrations, and water-storage capacity of a sandy soil under field conditions. Similarly, Agegnehu et al [18] found that, application of compost and biochar improved soil water and nutrient retention as well as water and nutrients uptake by the plants. In their study, however, little or no synergistic effect was observed. Abudjabha et al. [19] reported that application of biochar and compost enhanced microbial abundance due to enhanced formation of macropores and bioturbation. Biochar co-composting has been reported to minimize C [20] and N losses [21, 22], to decrease N_2_O [23] and CH_4_ [24] emissions compared to composting without biochar but the high diversity of biochar feedstocks makes it quite difficult to draw general conclusions on biochar co-composting effects.

This study was aimed to examine the effect of compost and biochar, applied singly, in combination or as co-compost on basal soil respiration, and soil quality indicators such as soil pH; soil microbial carbon; cation exchange capacity; total organic carbon, total nitrogen and available nitrogen concentration. The study is underpinned by the hypothesis that carbon-rich biochar and nitrogen-rich compost, applied together, can complement each other to improve soil quality indicators such as soil pH, C and N contents, soil microbial biomass and minimize soil C loss via respiration.

## Materials and Methods

### Soil, compost, biochar and composted biochar

Surface (0-20 cm) soil collected from an arable field at the University of Cape Coast Teaching and Research farm in Ghana (5°07’N, 1°17’W) was used for the study. The soil sampling site is located in the humid, coastal savannah agro-ecological zone with a mean annual precipitation of 1400 mm and a mean temperature of 20°C. The soils, which are developed on sandstones, shales and conglomerates are classified as a Haplic Acrisol [25]. The soil samples were collected from 20 different spots across a one hectare land, previously cropped with maize without any fertilization and bulked to form a composite sample for the study. The soil was well-drained and had a sandy loam texture (18, 9 and 73% clay, silt, and sand, respectively); with a pH of 4.3, and electrical conductivity of 200 µS cm^-1^; 9.3 g kg^-1^ total organic carbon ; 0.73 g kg^-1^ total nitrogen, and total phosphorus, potassium and magnesium contents of < 0.4, 11.9, and 9.3 mg 100 g^-1^, respectively.

The corn cob and rice husk biochars were produced by slow pyrolysis at approximately 450 °C in a locally produced kiln under low oxygen conditions. The coconut husk biochar was pyrolysed in a muffle furnace at 300 and 700 °C, respectively. The composted biochar was produced during the *UrbanFood plus* project in Tamale, Northern Ghana. The basic mixture of the treatments was poultry manure (15 vol-%; 68.7 kg m^-2^) and rice straw (60 vol-%; 75.4 kg m^-2^) and either corn cob or rice husk biochar (25 vol-%) (Volker Haring, personal communication).

The chemical properties of the soil and amendments used in the study are presented in Table 1.

**Table 1.** Chemical properties of amendments used in the study.

| Amendment | pH | TOC (%) | TN (%) | VOM | Ash (%) |
| --- | --- | --- | --- | --- | --- |
| Rice husk biochar | 7.2 | 44.26 | 0.75 | 25.7 | 20.1 |
| Corn cob biochar | 8.2 | 79.82 | 0.87 | 20.7 | 19.6 |
| Coconut husk biochar (300 °C) | 8.1 | 55.2 | 0.91 | 40.3 | 22.8 |
| Coconut husk biochar (700 °C) | 8.4 | 59.8 | 0.87 | 31.5 | 27.3 |
| Composted rice husk biochar | 8.5 | 43.7 | 0.92 | 25.2 | 29.6 |
| Composted corn cob biochar | 8.2 | 47.5 | 0.96 | 32.8 | 28.3 |
| Rice husk compost | 8.0 | 31.9 | 1.1 | 41.5 | 20.1 |

TOC = total organic carbon, TN = total nitrogen, VOM = Volatile organic matter.

### Experimental setup

A 30-day incubation was done with the surface (0-20 cm) Haplic acrisol soil samples with Kilner jars (500 ml) filled with 120 g air dry soils that had been sieved by mesh size of 2 mm. The soils were pre-incubated at 60% WHC and 25 °C for 10 days prior to addition of the amendments to re-initiate microbial activity after storage, and to minimise fluctuations in soil water content at the start of the experiment. At the end of the pre-incubation, each 120 g soil sample was re-adjusted to 60% WHC and mixed thoroughly with the soil amendments. The quantity of soil was adjusted so that upon addition of the amendments the mass of the soil-amendment mixture remained 120g. Each treatment was replicated 3 times for basal respiration measurements on days 7, 14, 21 and 30, respectively. The soils were destructively sampled on day 30 for MBC, pH, TOC, TON and CEC analyses. The different treatments and the respective quantities of amendments mixed with the soil during incubation are summarized in Table 2.

**Table 2.**
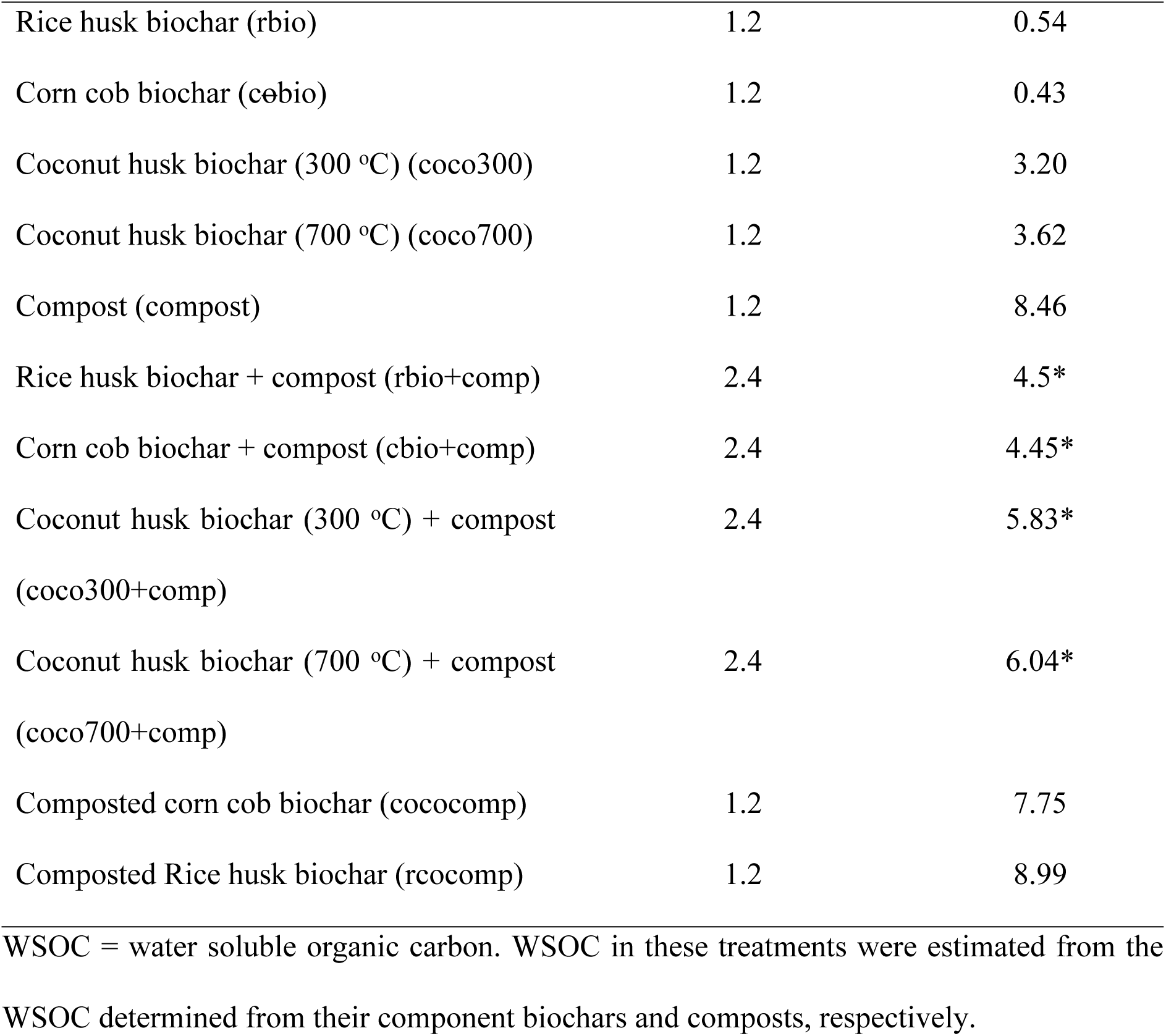
Treatment details (amendments-soil mixtures).

### Laboratory analyses

Total C and total N were analyzed by dry combustion (Vario EL Elementar Analysesysteme GmbH, Hanau, Germany) after grinding the dried samples. Soil pH was determined with a glass electrode (WTW 192, Ingold) in 0.01 M CaCl_2_ (1:25 w/v). The basal respiration was estimated weekly by titrating 10 ml 0.5 M KOH solution placed inside the beakers with 0.5M HCl. Microbial biomass C (Cmic) was determined by the chloroform fumigation extraction method [26]. 1.2g of incubated moist compost was fumigated at room temperature with ethanol-free CHCl_3_ for 24 hours in a desiccator. The fumigated sample and non-fumigated controls were extracted with 24ml of 0.05M K_2_SO_4_ by 30 min horizontal shaking and subsequent filtration (Whatman GF/A filters). The supernatant was subject to C and N determination with the Dimatoc 2000 analyzer. Cmic was calculated as difference between total C extracted from fumigated and non-fumigated treatments, divided by 0.45 [26].

Metabolic quotient (qCO2) was calculated as ratio of basal respiration and microbial biomass over the first week of incubation to describe the CO_2_-C produced per unit Cmic [27]. Further, the Cmic/ total C ratios were calculated to describe the C availability for microorganisms, respectively. All results were reported for oven dry (105°C). Hot water soluble C contents in the amendments used in the study were determined according to [28]. A mixture of 1.2 g of each amendment and 24 ml H_2_O were heated for 16 hours in a water bath at 80°C. The mixture was then cooled and shaken for 30 minutes prior to centrifugation. Then the supernatant was passed through a 0.45 μm filter (Whatman GF/A) and DOC in each mixture was analyzed with a Dimatoc 2000 automatic analyzer (Dimatec, Essen, Germany).

The exchangeable cations and CEC were determined using the ammonium chloride (at pH 7) extraction method. About 2.5 g of the sample was weighed into a 50 ml beaker and 10ml of the exchange solution was added. The beaker was covered with watch glass and allowed to stand for 24 hours. Percolation was done after 24 hours by rinsing the sample with the solution into the filter paper and collected in 100 ml measuring flask. The percolation lasted about 4 hours with approximately 90 ml of the percolate in the flask. The concentration of the exchangeable cation was measured with the Inductively Coupled Plasma Spectroscopy (ICP-OES). The CEC was calculated as the sum of the exchangeable cations (Ca^2+^, Mg^2+^, Na^+^ and K^+^).

### Statistical analyses

The statistical analyses were done with the 12^th^ edition of GenStat statistical software. Descriptive analyses, and tests for normality and homogeneity of variances were conducted prior to a one-way analyses of variances (ANOVA). Analysis of variance (ANOVA) was used to test the effects of treatments on soil quality parameters. Pearson’s correlation analysis was used to determine whether there were significant interrelationships among the measured properties of the soils. Treatment means which were found to be significantly different from each other were separated by Duncan tests.

## Results

### Basal respiration

The effects of biochar and / or compost application on basal respiration are presented in Figs 1a and 1b.

**Fig 1a.**
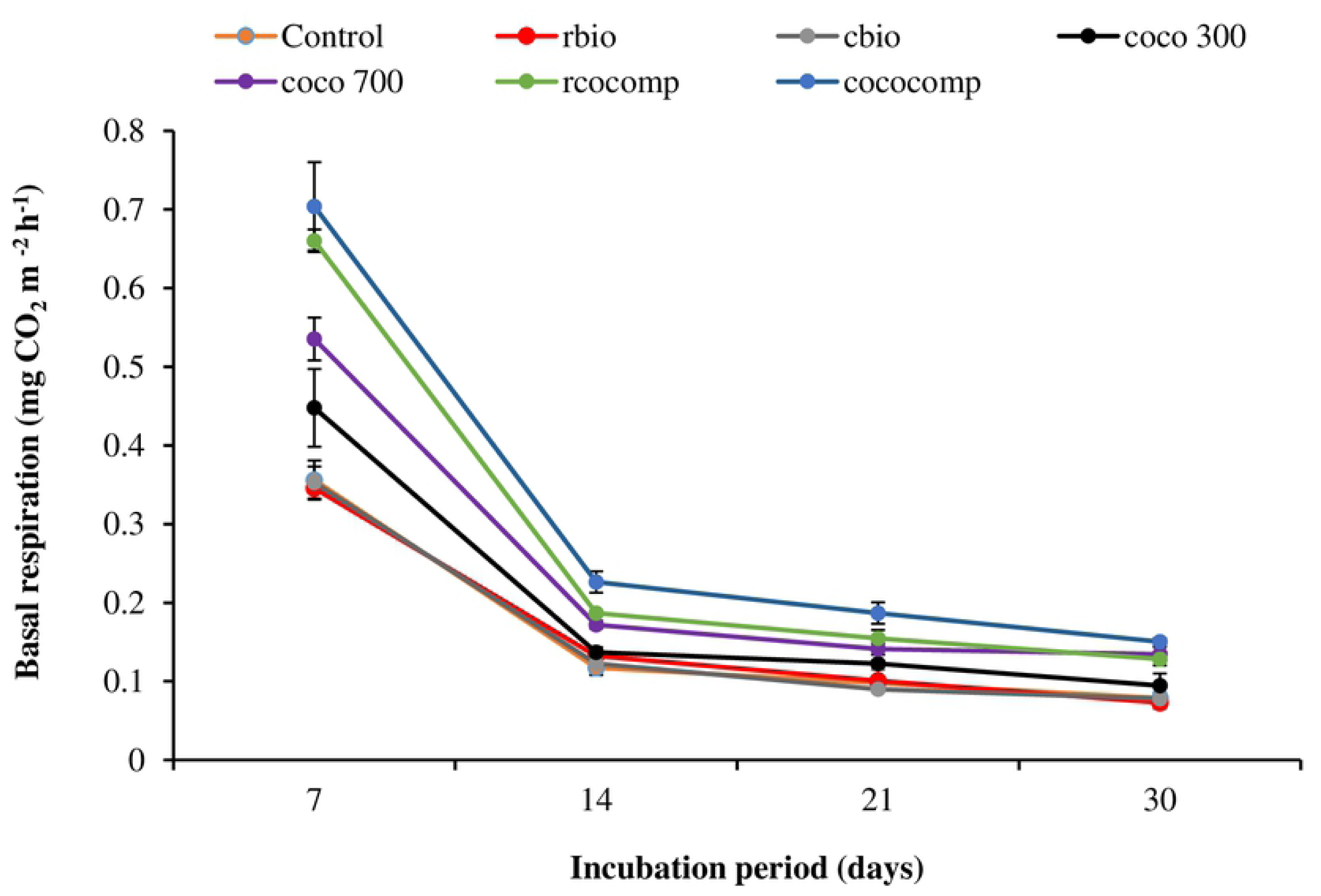
Basal respiration rate following the application of biochar or composted biochar to soil.

**Fig 1b.**
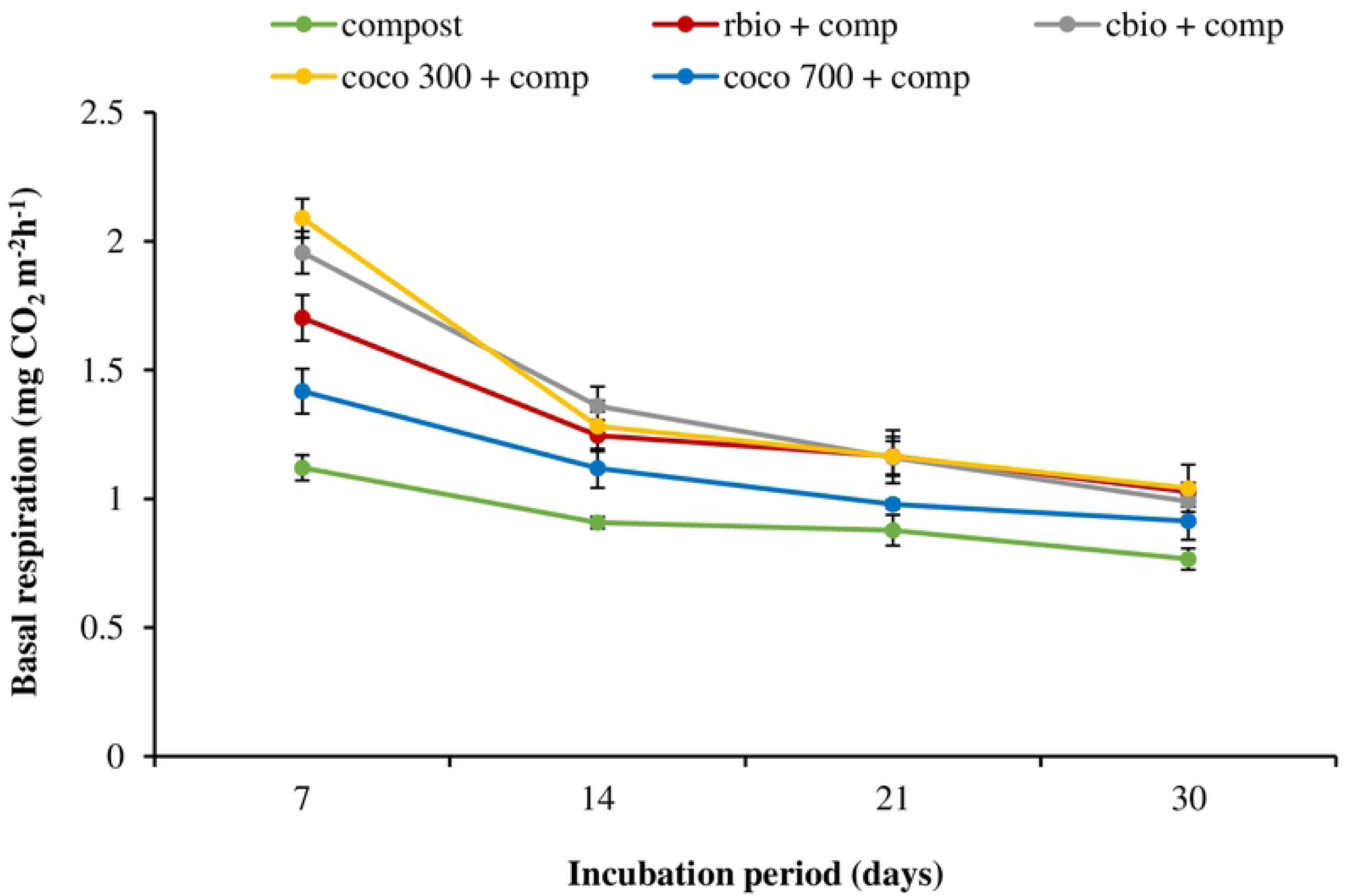
Basal respiration rate following the application of compost only or combined compost and biochar to soil.

Basal respiration peaked on day 7 in all the treatments (Figs 1a and 1b), with all the amended soil except rbio and cbio showing higher peak basal respiration than the control. Basal respiration in the combined biochar and compost treatment were relatively higher than their corresponding sole biochar treatments. Sole addition of compost resulted in higher basal respiration than application of biochar co-compost. Basal respiration recorded from day 14 to day 30 were similar in all the sole biochar treatments.

### Effect of sole biochar, combined biochar and compost or composted biochar applications on soil properties

Tables 3 shows the effects of the amendments: sole biochar, sole compost, combined compost and biochar or composted biochar on soil pH, total organic carbon (TOC), total nitrogen (TN) contents and microbial biomass carbon (MBC) while Table 4 shows the amendments on soil total respiration (TR), MBC/TOC, Metabolic quotient (qCO2) and TOC/TR after 30 days of incubation in the control and amended soils at the end of the incubation period.

**Table 3:** Effects of sole biochar, sole compost, combined compost and biochar or composted biochar on soil pH, TOC, TN and MBC after 30 days of incubation.

| Treatments | pH | TOC (%) | TN (%) | MBC (µg/g) |
| --- | --- | --- | --- | --- |
| **Control** | 4.39 | 0.81 ± 0.03a | 0.10 ± 0.00a | 62.71 ± 16.53a |
| **rbio** | 4.67 | 1.59 ± 0.56cde | 0.14 ± 0.05b | 62.96 ± 18.31a |
| **cbio** | 4.80 | 1.55 ± 0.09cde | 0.11 ± 0.00ab | 68.23 ± 8.33a |
| **coco300** | 5.02 | 1.40 ± 0.02bcd | 0.11 ± 0.00ab | 72.62 ± 2.27a |
| **coco700** | 5.47 | 1.52 ± 0.02cde | 0.10 ± 0.00a | 73.98 ± 3.26a |
| **compost** | 6.38 | 1.11 ± 0.05b | 0.12 ± 0.00ab | 217.63 ± 55.09d |
| **rcocomp** | 5.64 | 1.16 ± 0.03b | 0.11 ± 0.00ab | 106.50 ± 9.40ab |
| **cococomp** | 5.75 | 1.35 ± 0.04bc | 0.11 ± 0.00ab | 109.25 ± 6.72ab |
| **rbio + comp** | 5.98 | 1.47 ± 0.02cde | 0.12 ± 0.00ab | 145.64 ± 78.87bc |
| **cbio + comp** | 6.09 | 1.76 ± 0.04e | 0.12 ± 0.00ab | 209.47 ± 16.79d |
| **Coco 300 + comp** | 6.20 | 1.61 ± 0.02cde | 0.13 ± 0.00ab | 196.06 ± 23.09cd |
| **coco 700 + comp** | 6.68 | 1.70 ± 0.02de | 0.12 ± 0.00ab | 185.22 ± 33.17cd |
| ***p* value** |  | <0.001 | 0.207 | < 0.001 |
| **LSD (0.05)** |  | 0.279 | 0.025 | 53.291 |

**Table 4:** Effects of sole biochar, sole compost, combined compost and biochar or composted biochar on soil total respiration (TR), MBC/TOC, Metabolic quotient (qCO2) and TOC/TR after 30 days of incubation.

| Treatments | MBC/TOC | TR<br>(mg CO<sub>2</sub> (m<sup>2</sup> h<sup>-1</sup>)) | MBC/TR<br>(qCO<sub>2</sub>) | TOC/TR |
| --- | --- | --- | --- | --- |
| **Control** | 77.48 ± 23.34abc | 0.65 ± 0.03a | 95.93 ± 21.17de | 1.25 ± 0.09bc |
| **rbio** | 39.60 ± 22.86a | 0.65 ± 0.02a | 96.51 ± 27.59de | 2.45 ± 0.92e |
| **cbio** | 44.07 ± 7.11a | 0.64 ± 0.02a | 106.20 ± 14.62e | 2.40 ± 0.07e |
| **coco300** | 51.72 ± 2.25ab | 0.80 ± 0.07ab | 91.12 ± 9.93de | 1.76 ± 0.13d |
| **coco700** | 48.52 ± 1.93ab | 0.98 ± 0.04bc | 75.29 ± 2.77cd | 1.55 ± 0.04cd |
| **compost** | 195.59 ± 46.88d | 3.67 ± 0.14e | 59.00 ± 13.49bc | 0.30 ± 0.01a |
| **rcocomp** | 92.17 ± 8.97bc | 1.13 ± 0.03cd | 94.45 ± 11.08de | 1.02 ± 0.04b |
| **cococomp** | 81.15 ± 7.29abc | 1.27 ± 0.07d | 86.37 ± 7.22de | 1.07 ± 0.08b |
| **rbio + comp** | 98.83 ± 54.24c | 5.14 ± 0.06g | 28.32 ± 15.32a | 0.29 ± 0.00a |
| **cbio + comp** | 118.75 ± 11.66c | 5.46 ± 0.12h | 38.30 ± 2.29ab | 0.32 ± 0.01a |
| **coco300+comp** | 121.70 ± 13.22c | 5.57 ± 0.35h | 35.39 ± 6.11ab | 0.29 ± 0.02a |
| **coco700+comp** | 108.85 ± 20.52c | 4.43 ± 0.30f | 41.72 ± 5.96ab | 0.39 ± 0.03a |
| ***p* value** | < 0.001 | < 0.001 | < 0.001 | < 0.001 |
| **LSD (0.05)** | 41.080 | 0.246 | 22.815 | 0.456 |

### Soil pH

All the amended soils showed higher soil pH than the control (Table 3). Addition of biochar and / or compost or composted biochar increased soil pH by 0.28 - 2.29 with 700 + comp treatment recording the highest increase in soil pH. The sole compost and composted biochar treatments increased soil pH more than the sole biochar treatment. The combined biochar and compost treatments showed higher soil pH values than their corresponding sole biochar treatments and composted biochar amended soils.

### Total organic carbon (TOC) and total nitrogen (TN)

Total organic carbon in all the amended soils were greater than the control (Table 3). TOC in the sole biochar and combined biochar and compost treatments were greater than their corresponding sole compost and composted biochar treatments. However, regardless of the feedstock, addition of biochar together with compost did not increase TOC more than the sole application of biochar. TN found in all the treatments were not statistically different from the control except for rbio which increased TN by 40%.

### Microbial biomass carbon (MBC)

MBC in all the sole biochar and composted biochar treatments were similar to the control. However, when biochar and compost were applied together, MBC was increased by 82.9 to 146.8µg g^-1^ compared to the control. Also, sole application of compost increased MBC more than the corresponding composted biochars.

### MBC / TOC

The ratio of MBC to TOC in all the treatments were similar to the control except for the sole compost treatment. Applying compost alone increased MBC/TOC by 152.4%. Although applying biochar together with compost increased MBC/TOC more than their corresponding sole biochar treatments, differences between them and the control were not statistically significant.

### Total respiration (TR)

Total respiration was calculated by summing up the basal respiration measured on days 7, 14, 21 and 30 for each treatment. The sole biochar treatments showed no significant differences in TR as compared to the control treatment except for coco 700 (Table 2). TR in the amended soils followed the order combined biochar and compost > sole compost > composted biochar > sole biochar > control. Among the combined biochar and compost treatments, coco300 + comp gave the highest TR value of 5.57 mg CO_2_ m^-2^ hr^-1^ which was also the highest among all the treatments.

### Metabolic quotient (qCO2) (MBC/ basal respiration)

Microbial quotient (qCO_2_) was lower in the sole compost and the combined biochar and compost treatments compared to the control. However, the sole biochar and the composted biochar treatments were not statistically different from the control.

### Cation exchange capacity (CEC)

CEC in all the treatments were significantly higher relative to the control (Fig 2). The application of biochar together with compost increased CEC more than biochar applied alone. CEC in the composted biochar treatments were similar to those in the sole compost treatments but these were higher than those found in their corresponding combined compost and biochar treatments.

**Fig 2.**
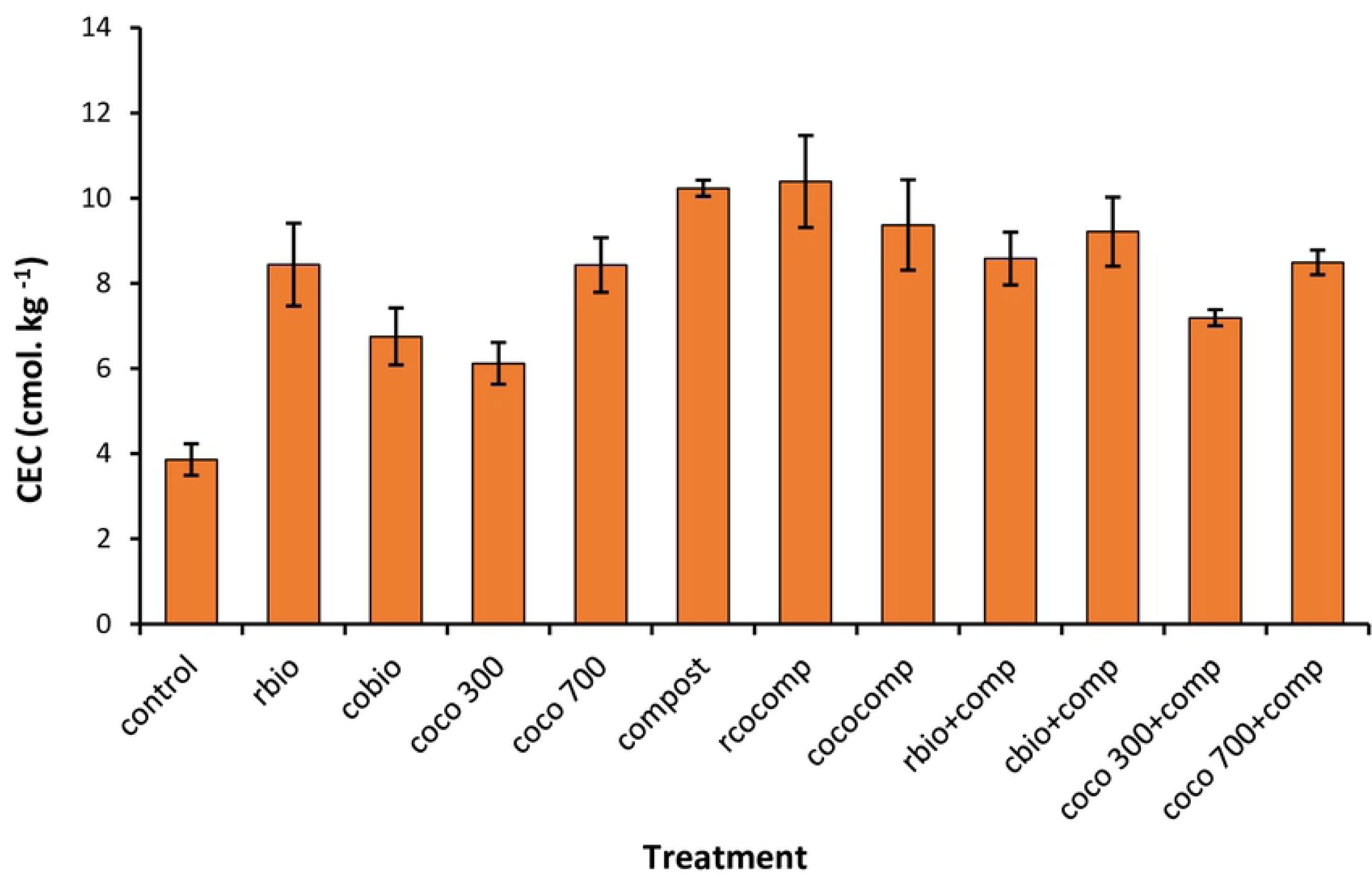
Effect of biochar and / or compost application on CEC.

### Ammonium nitrogen (NH_4_ ^+^-N)

Inorganic NH_4_ ^+^-N concentrations in all the amendments were lower (P < 0.05) than the control except for rbio and cbio (Fig. 3).

**Fig 3.**
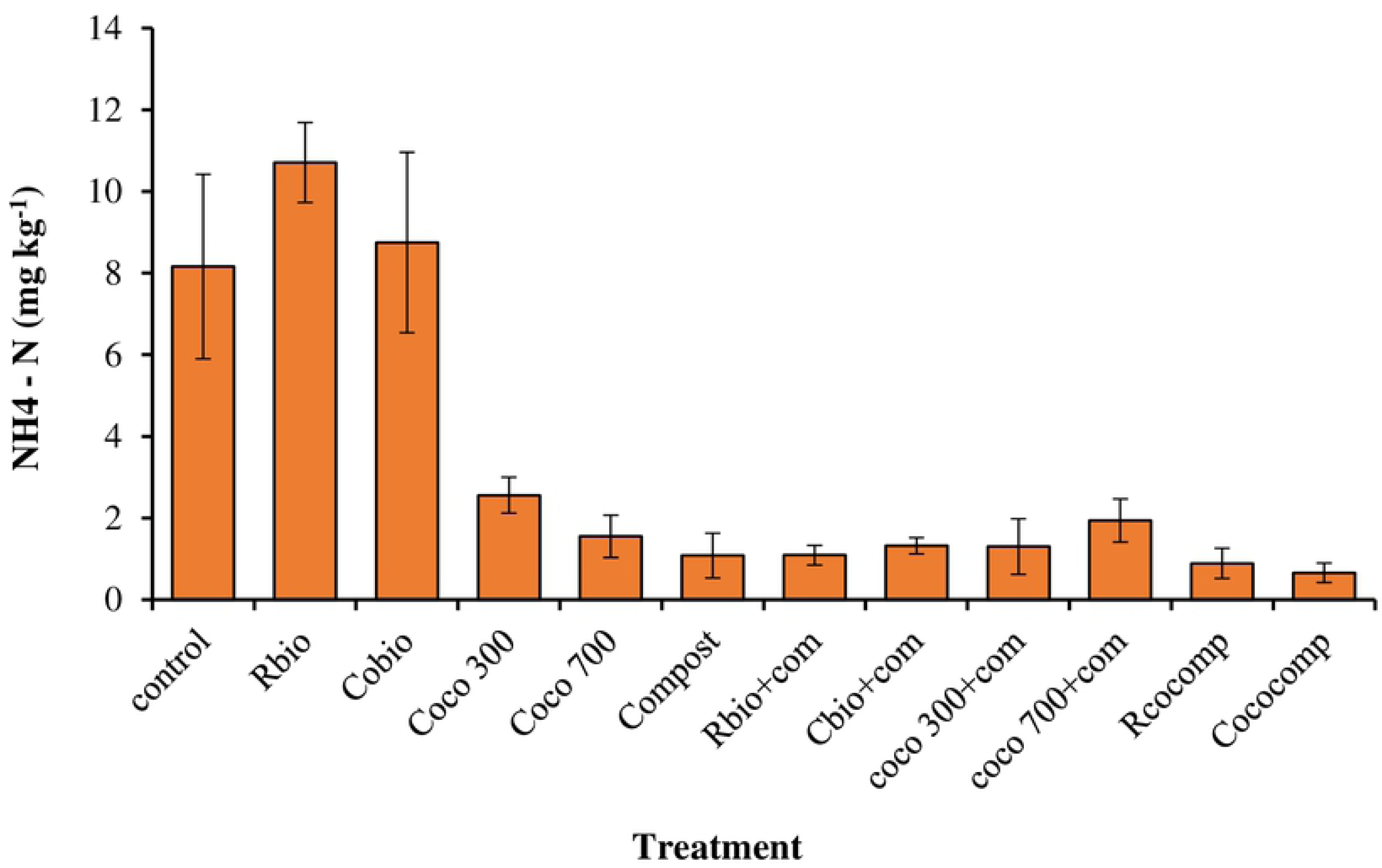
Effect of biochar and/ or compost application on NH_4_ ^+^ – N.

At the end of the incubation period, the NH_4_ ^+^-N concentrations in the sole biochar amended soil were higher than their corresponding combined biochar and compost treatments. No significant differences were however observed among the sole compost, composted biochar and the combined compost and biochar treatments.

### Nitrate nitrogen (NO_3_ ^-^-N)

Inorganic NO_3_ ^-^-N concentrations were higher in the sole compost, composted biochar and the sole biochar treatments as compared to the control (Fig. 4).

**Fig 4.**
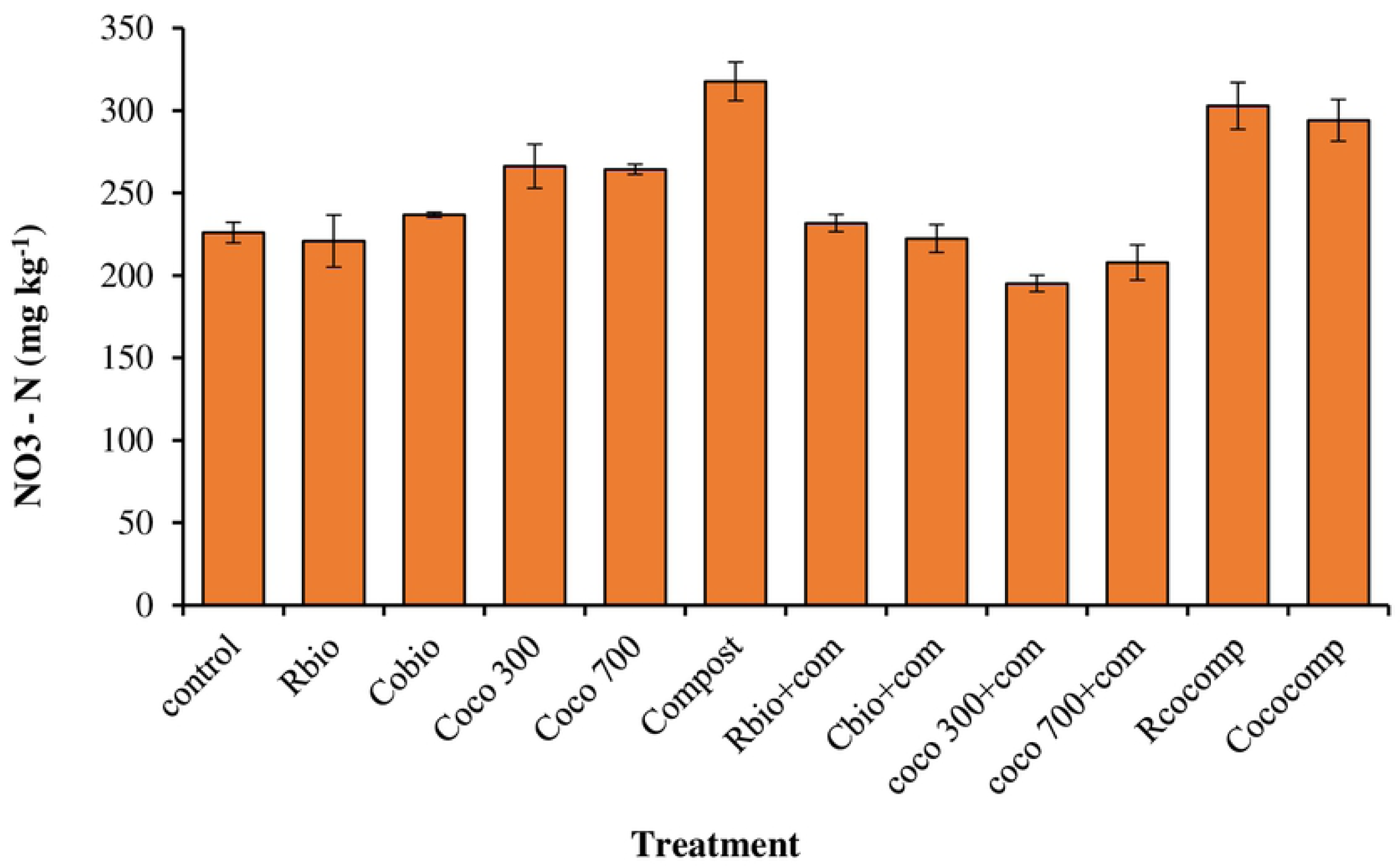
Effect of biochar and/ or compost application on NO_3_ – N.

The NO_3_ ^-^-N concentrations in the sole biochar amended soil were higher than their corresponding combined biochar and compost treatments. The highest NO_3_ ^-^-N concentrations was recorded in the sole compost treatment followed by the composted biochar treatments.

### Biochar and compost mineralisation (% SOC)

C mineralized in biochar and / or compost amended soils are presented in Figs 5 and 6, respectively.

**Fig 5.**
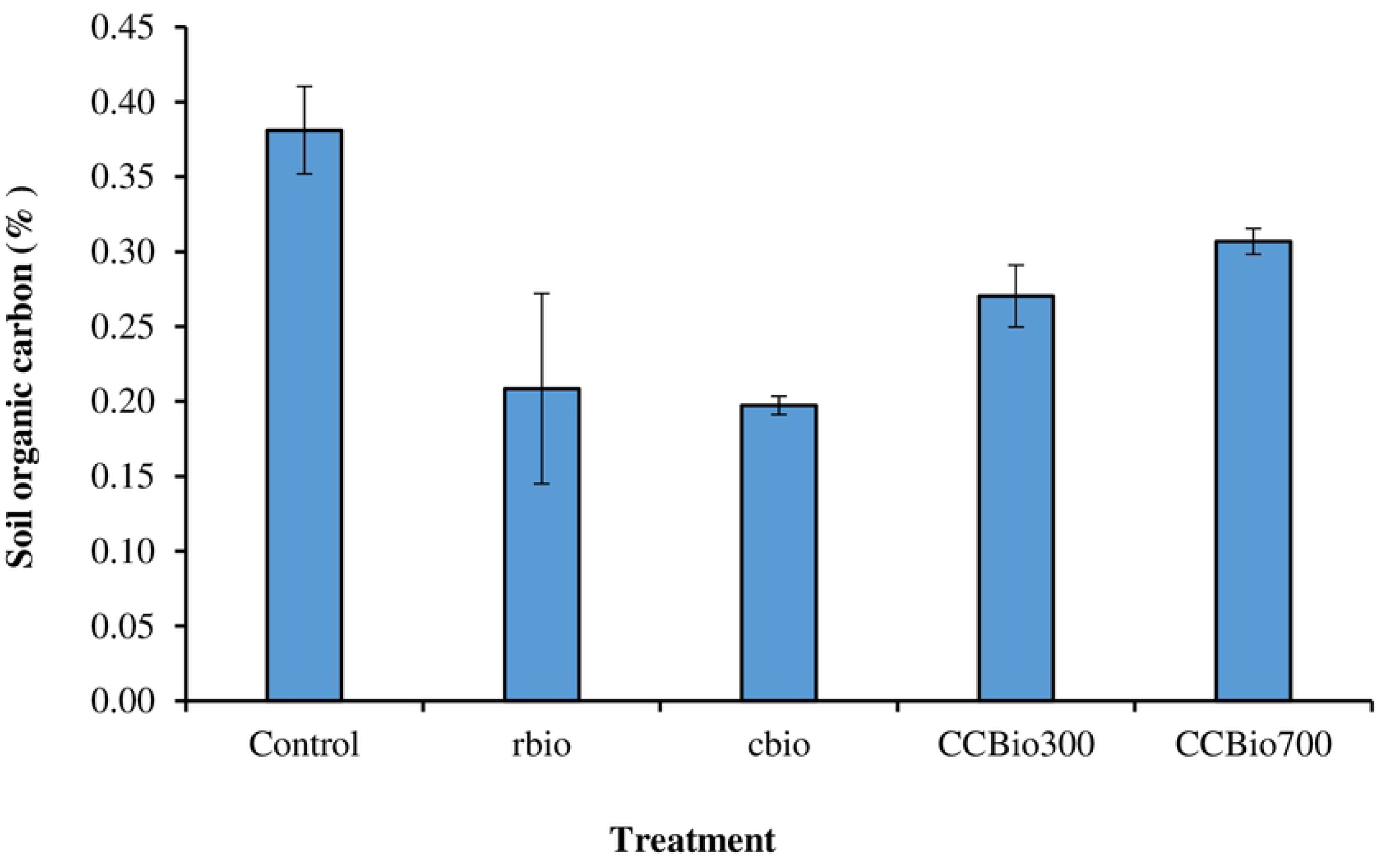
Effect of biochar and / or compost application on biochar mineralization.

**Fig 6.**
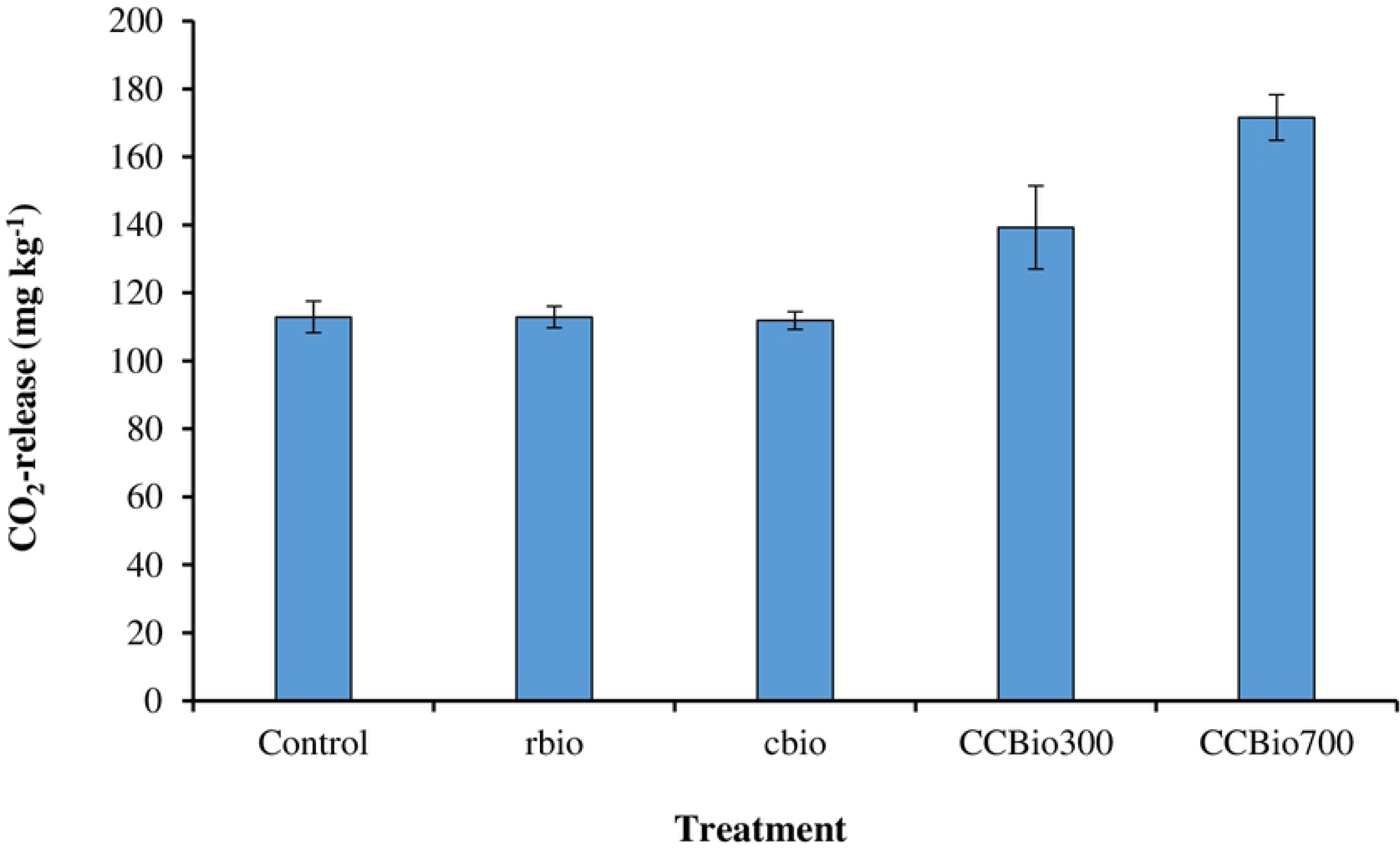
Effect of biochar and/ or compost application on compost mineralization.

### CO_2_ release biochar and compost-biochar mixtures treatments

CO_2_ release following biochar and / or compost application are shown in Figs 7 and 8, respectively.

**Fig 7.**
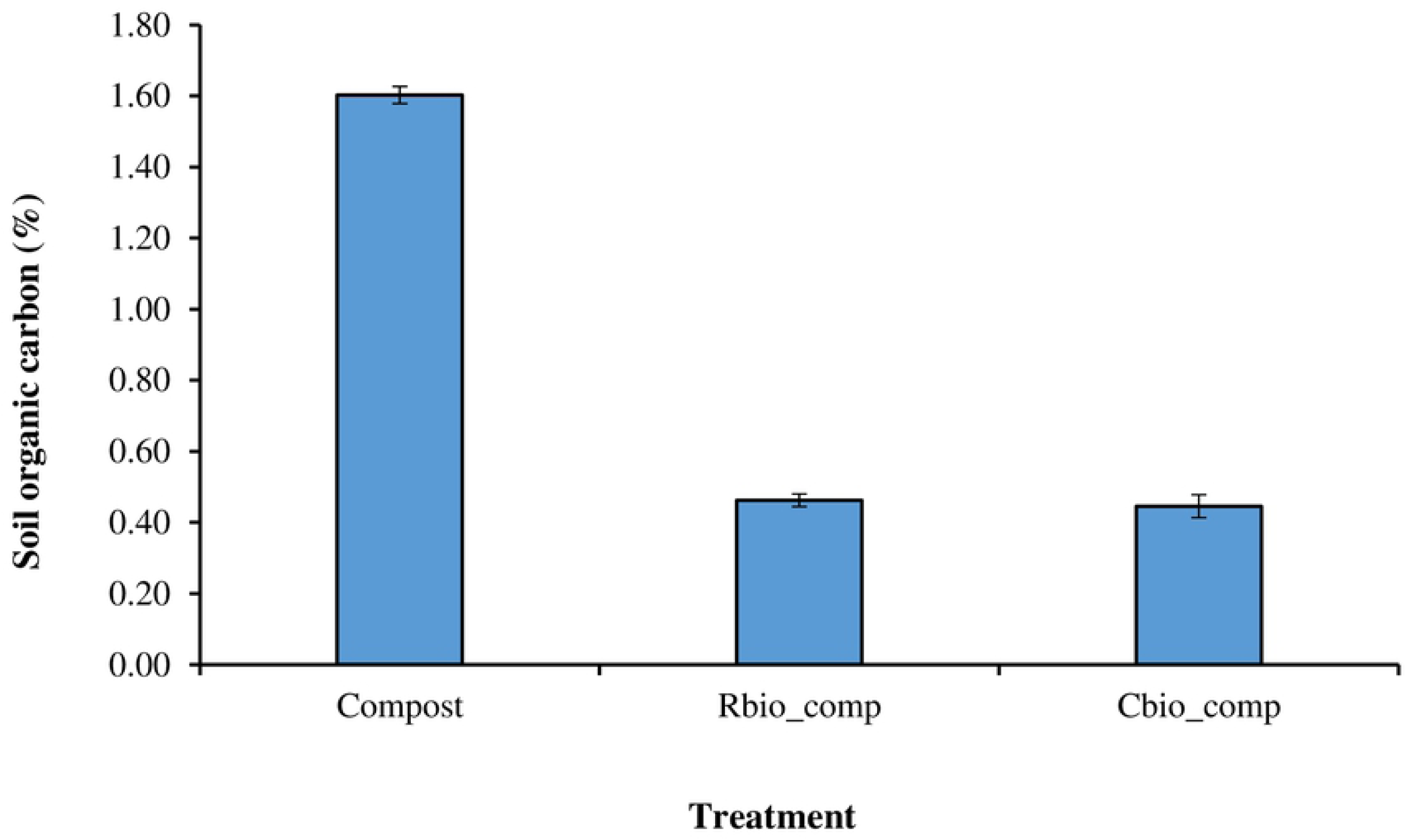
Effect of biochar and/ or compost application on CO_2_ release from biochar treatments.

**Fig 8.**
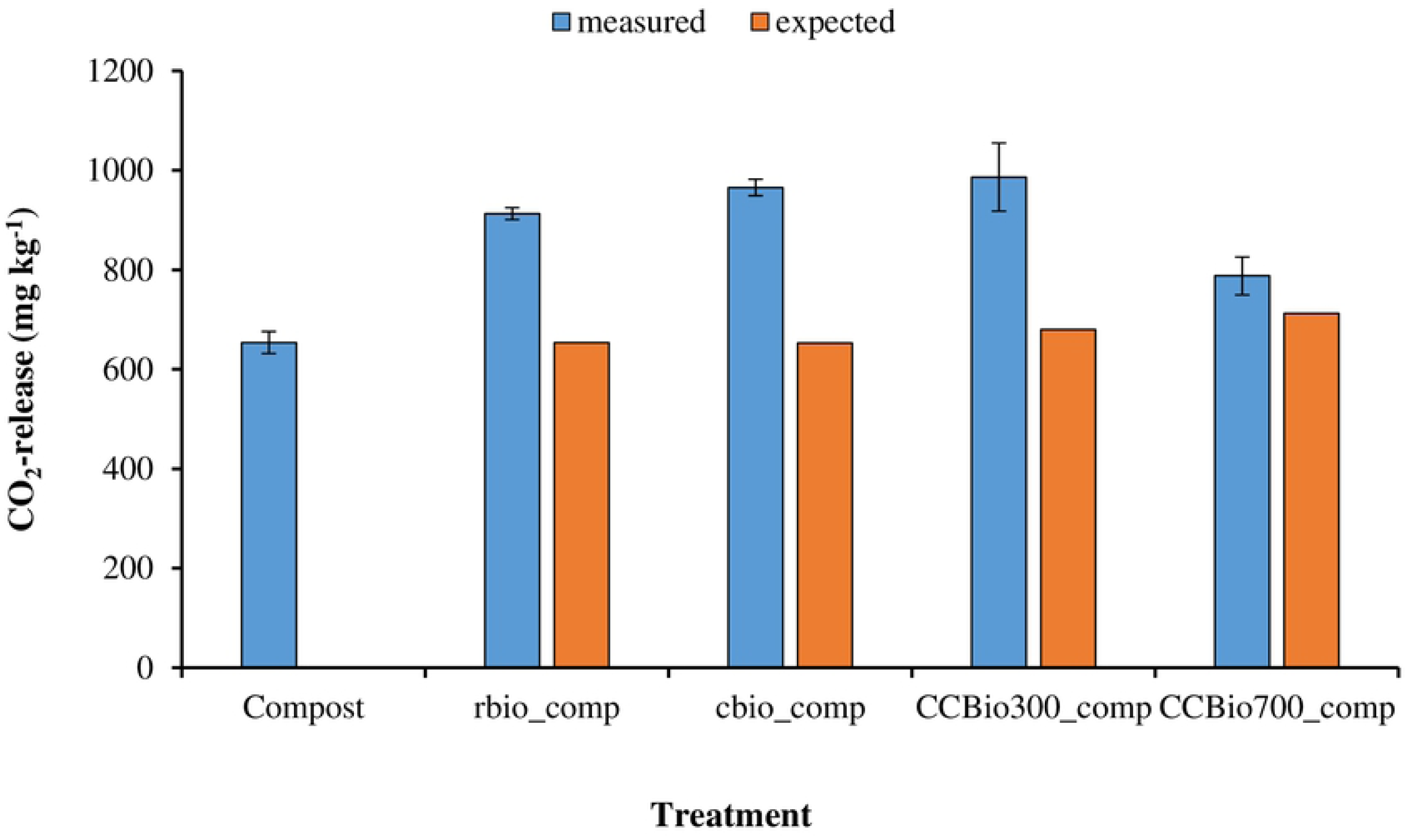
Effect of biochar and/ or compost application on CO_2_ release from compost-biochar mixtures.

## Discussion

Results from the study showed that sole biochar additions generally, did not significantly increase basal respiration compared with the control. This finding is in good agreement with Zavalloni et al. [29] who found that respiration rates in soil treated with coppiced woodland-derived biochar were not significantly different from control soil. Previous authors [30,31] opined that low CO_2_ emissions from biochar amended soils can be attributed to chemisorptions of the respired CO_2_ on the biochar surface. Basal respiration from the combined biochar and compost, sole compost and composted biochar treatments, respectively, were greater than the sole biochar treatments (Figure 1a and 1b). Basal respiration is an indication of microbial activity in soil. The higher basal respiration found in the sole compost, combined biochar and compost and the composted biochar treatments is an indication of increased biological activity in those treatments. Additionally, the study showed rather unexpectedly that mixing compost with very inert biochars (Rbio and Cbio) greatly increased mineralization, with the lowest effects occurring rather with the most labile biochar (coco700). The underlying reason for this observation is not clear but it raises the critical questions whether the biochar removed some toxic compounds or provided additional nutrients, which would have stimulated the mineralization process. Further studies need to be done to establish the mechanisms underpinning this observation.

Water soluble organic carbon (WSOC) content in the compost treatment was greater than in the corn cob biochar, rice husk biochar and the coconut husk biochars, respectively (Table 2). WSOC contents in the combined biochar and compost amendments were higher than the sole compost amendment. This is indicative that most of the labile C that drove microbial activity would have been released from both the biochar and the compost in the mixes. In agreement with Méndez at al. [30], we suggest that, here, differences in basal respiration from the compost and / or biochar amended soils can be related to the varying labile carbon content of the amendments. Benito et al. [32] reported that, when immature compost is applied, its high content of water soluble carbon can lead to stimulation of microbial activity followed by an increased carbon dioxide (CO_2_) fluxes and higher soil organic matter (SOM) decomposition through priming effect. The study showed that coconut husk biochars were more labile than the corn cob and rice husk biochars, respectively. Moreover, although the coconut biochar pyrolysed at 700°C appeared to be more labile than the one pyrolysed at 300°C, this did not reflect in a higher MBC. This suggest that perhaps a greater concentration of some secondary metabolites or other components in the former suppressed microbial growth compared with the latter, but this requires further investigation. The results also revealed that co-composting greatly reduces the labiality of the product, which also reflected in the MBC. Unfortunately, we did not estimate the proportion of MBC remaining in the co-compost.

In this study, MBC in sole biochar amended soils were significantly lower compared to those in the sole compost and the combined compost and biochar. However, there was no difference between MBC from sole biochar and composted biochar treatments. The results is in accordance with Liu et al. [33] who observed no effect of corn cob straw biochar on MBC in a laboratory experiment in which corn cob pyrolysed at 550°C had been incorporated in a sandy loam at 15t ha^-1^. Méndez et al. [30] and Thies and Rillig [31] argued that the electrical conductivity, metal and phenolic substances of biochar can have a negative effect on soil microbial activity, reducing the respired CO_2_. In contrast to our results, Khan et al [34] observed that many studies reported increases of MBC with biochar additions. According to Kuzyakov et al. [35], a small fraction of labile C in biochar can be mineralized within a short period after its application and can stimulate microbial efficiency in soils [36]. Thus, biochar can supply C substrate for soil microbes [37], thereby enhancing the activities and efficiency of microbes in soils. This microbial growth is believed to induce soil C mineralization or degradation [38]. In this study, application of biochar alone did not show any significant increase in MBC. Our findings is in agreement with Jindo et al. [39] who reported lower MBC for biochar than non-biochar treatments and related it to low labile C contents in biochar amended soils.

According to Chen et al. [40], soil microbial biomass greatly depends on soil organic matter as substrate and so a decrease in SOC is likely to result in low soil microbial biomass [40]. Thus, the higher MBC in the sole compost and the combined compost and biochar amended soils are attributable to their greater organic C contents [41]. This observation is further confirmed by the positive correlations between soil microbial biomass and total organic carbon contents (r^2^ = 0.202, *P* = 0.237) found in this study. Farrel et al. [42] reported that MBC is a useful indicator of soil C stabilization following soil management and Bierdeman and Harpole [43] posited that increases in MBC suggest higher availability of labile C inputs. In our study, the trends in MBC contents in the differently amended soils was similar to those observed for the basal respiration values. For instance both low MBC and the low basal respiration in the sole biochar treatments indicate low microbial activity. Further, MBC in the combined compost and rice husk or corn cob biochar treatments were greater than their corresponding composted biochars, suggesting that decomposable C may have been stabilized in the composted biochars. In other words, the MBC/SOC ratio shows substrate availability and the portion of total soil carbon immobilized in microbial cells.

In this study, the relatively lower MBC/SOC ratio in the sole biochar amended soils may be attributed to the inhibition of microbial immobilization, whereas a relatively higher MBC/SOC ratio in the sole compost may imply stimulation of microbial immobilization. Combined compost and biochar and composted biochar treatments possibly contained more and diversified organic C substrate, leading to a higher carbon availability. According to Sparling [44], soil MBC/SOC ratio, or microbial quotient, has been widely used as an indicator of future changes in organic matter due to modifications of soil conditions. Furthermore, Joergensen and Scheu [45] mentioned that this ratio is used as a basis for comparison of soil quality parameter across soils with different organic matter contents. Thus, the MBC/SOC ratio mirrors the relationship and interactions between the MBC and mineral SOC [46]. The large amounts of recalcitrant carbon in biochar amended soils would have resulted in lower rate of decomposition and consequently slower transformation of particulate organic matter into mineral soil component [47]. This is evidenced by lower TON and basal respiration in the sole biochar amended soils.

Previous studies reported positive correlations between litter layer C/N ratio and respiration [48,49]. In this study, a positive correlation was found between soil C/N ratio and peak respiration at day 7 but this was not statistically significant. The findings go in line with a positive correlation between qCO_2_ and the soil C-to-nutrient ratios that Spohn and Chodak found in beech, spruce, and mixed forests. Three reasons have been proffered for these observed relationships. First, Craine et al. [51] posited that microorganisms burn readily available C in order to gain energy to acquire N from more recalcitrant forms of organic matter [51] or in order to have physical access to the N incorporated in organic compounds. However, the capacity of microorganisms in N-limited soils, where the payoff in terms of N, is very small to invest N into the production of exoenzymes and release them to acquire C, have been challenged.

Secondly, Russel and Cook et al. [52] and Manzoni et al., [53] attributed the relationship to overflow respiration, which implies that microorganisms uncouple respiration from energy production and only respire C to dispose it off. However, the relevance of microbial overflow respiration in ecosystems has been questioned [54] on the basis that first, in order to dispose off C via respiration, N is required to maintain the proteins of the respiratory chain [54]. Second, microorganisms may use the C that is in excess of that required for growth of cells for C storage, buildup of structural defences, viral repellents or establishment of symbiosis. However, all these physiological processes also require N and that amount of C that microbes can store or invest in the above-named processes is limited. Finally, Carreiro et al. [55]; Michel and Matzner [49] and Gallo et al., [56] explained that activity of oxidative enzymes involved in the degradation of aromatic compounds, decreases with increasing N concentration. Thus, decreased lignolytic activity might decrease microbial respiration in litter with low C:N ratios [55,57].

Previous studies have suggested that the higher the qCO_2_ in a treatment, the more C is channelled into the energy metabolism and the lower the efficiency of microbial biomass C formation. Some authors have argued that low qCO_2_ rates indicates high microbial efficiency, which implies that organic matter decomposition is higher and that C mineralization is more stimulated in less stressed ecosystem due to abundance and higher activity of soil microbes. In contrast, in this study, the efficiency of microbial C formation was relatively lower in the sole compost treatment than in the combined biochar and compost treatments. The sole biochar amendments showed lower water soluble C than the sole compost. Therefore, it was not unexpected that microbial biomass C and basal respiration were both greater in the sole compost treatment than in the sole biochar treatments. However, it seems that basal respiration was disproportionately greater than MBC in the sole compost amended soils, hence the low qCO_2_ measured in the sole compost treatments.

The results showed that biochar and/or compost additions increased soil pH above that of the control soil. The influence of biochar additions on soil pH has been reported severally. Méndez et al. [58] observed a pH increment on a Haplic Cambisol after the addition of sewage sludge-derived biochar. Kloss et al. [59] also found slight increment of soil pH (0.3 units) in an acid soil after application of woodchip-derived biochar, whereas Jien and Wang [60] observed a significant increase (from 3.9 to 5.1) in an Ultisol following addition of wild lead tree, and waste wood biochar. It must be pointed out that pH of 6.6 to 7.3 is favorable for microbial activities that contribute to the availability of nitrogen, sulfur, and phosphorus in soils [61], and a pH value exceeding 10 can have negative effects on soil nutrient availability, but Schmidt et al. [62] concluded that only the application of larger amounts of biochar will lead to large changes in a soil’s pH value.

In this study, TOC in all the amended soils were greater than the control. Previous studies have indicated that total SOC was significantly increased due to the applications of various biochar types [63,64,65]. Here, combined biochar and compost treatments increased TOC by 81.5-117.3% compared to the control while the composted rice husk and corn cob biochars increased TOC by 43.2 and 66.7 % respectively. The significantly higher TOC observed in the combined biochar and compost treatments reflected the amounts of amendment C originally added to the soil during incubation in that 1.2g each of the compost and biochar added to the soils of the combined additions of biochar and compost treatments. Sole compost application increased TOC by only 37% compared to the control. This suggests that combined application of biochar and compost synergistically increased soil TOC content. This has positive implications for soil C sequestration. Kimetu and Lehmann [64] found that soils initially containing low C is more efficient in stabilizing SOC after biochar additions than soil initially richer in C. In this study, only one soil type was involved hence this observation could not have been verified.

Our results agrees with Xie et al. [65] who reported that biochar addition increased soil N contents. In contrast, Lehmann and Joseph [66] reported that pure biochar does not directly enrich the soil with nutrients. Again, Lehmann et al., [67] opined that biochar addition may have elevated the soil C/N ratio, thereby reducing N mineralization. In this study, sole biochar and sole compost application increased TON by 12 to 27%. TON in the combined biochar and compost treatments were greater than in the corresponding composted biochar treatments. Following the ideas of Schulz and Glaser [68] we hypothesize that composting biochar would have had stabilizing effects on early decomposable components of their compost, resulting in the relatively N contents in the composted biochar treatments than in the combined compost and biochar treatments. However, TON in the composted biochar treatments was higher than in the sole biochar treatments.

Berglund et al. [69] and Deluca et al. [70] explained that notwithstanding the generally, very low N content of biochar, net nitrification could be stimulated following biochar addition to soil particularly where soil pH is optimal. Thus, the greater TON contents in the combined biochar and compost treatments was not unexpected. However, the results in this study do not show evidence of any priming effect. In another study, Yao et al., [71] observed absent or even negative effect on nitrification, mainly depending on soil pH. In this study, none of the amendments increased soil pH to above 10 and so no net effect on nitrification could have occurred. With respect to CEC, biochars increased soil CEC, a result which disagrees with Méndez et al., [58]. However, CEC in sole biochar treatments were lower than in the sole compost, combined biochar and compost and the composted biochar treatments, which is indicative of the low inherent CEC of biochar compared to compost [72].

## Conclusions

The study has demonstrated that additions of compost and biochar, and composted biochar enhanced soil quality in terms of increased TOC, TON, MBC, CEC and soil pH. Furthermore, the study showed that biochar application resulted in soil C stabilization exemplified by lower of CO_2_ emissions in biochar amended soils compared to compost amended treatments attributable to a relatively higher labile C availability in the sole compost treatment. The efficiency of C conversion into MBC was enhanced in all compost treatments regardless of whether the compost was added solely or in combination with biochar.

## Acknowledgements

We acknowledge Ruhr University of Bochum for providing office and laboratory space during the 3-montth research fellowship during which the project was done; University of Cape Coast for the study leave to undertake the fellowship; Ms. Sabine Frolich, Ms. Katja Gonschorek and Ms. Heidrun Kerkhoff for their assistance in laboratory analysis, as well as Dr Volker Harring, Dr Stefanie Heinze and Dr. Blessing Chinyere Obika at the Department of Soil Science/Soil Ecology, Ruhr University of Bochum for the laboratory assistance and technical advice they offered.

## Supporting information

**S1 Table. Pearson correlation matrix. (**\*\*. Correlation is significant at the 0.01 level; *. Correlation is significant at the 0.05 level.

